# A gain and dynamic range independent index to quantify spillover spread to aid panel design in flow cytometry

**DOI:** 10.1101/2020.10.31.363150

**Authors:** Debajit Bhowmick, Frank van Diepen, Anita Pfauth, Renaud Tissier, Michelle Ratliff

**Author notes:** Correspondence to: Debajit Bhowmick.

## Abstract

In conventional flowcytometry one detector (primary) is dedicated for one fluorochrome. However, photons usually end up in other detectors too (fluorescence spillover). ‘Compensation’ is a process that corrects the spillover signal from all detectors except the primary detector. Post ‘compensation’, the photon counting error of spillover signals become evident as spreading of the data. The spreading induced by spillover impairs the ability to resolve stained cell population from the unstained one, potentially reducing or completely losing cell populations. For successful multi-color panel design, it is important to know the expected spillover to maximize the data resolution. The Spillover Spreading Matrix (SSM) can be used to estimate the spread, but the outcome is dependent on detector sensitivity. Simply, the same single stained sample produces different spillover spread values when detector(s) sensitivity is altered. Many researchers mistakenly use this artifact to “reduce” the spread by decreasing detector sensitivity. This can result in diminished capacity to resolve dimly expressing cell populations. Here, we introduce SQI (Spread Quantification Index), that can quantify the spillover spread independent of detector sensitivity and independent of dynamic range. This allows users to compare spillover spread between instruments having different types of detectors, which is not possible using SSM.

## Introduction

Flow Cytometry is a powerful technique to monitor and study huge number of cells within a few minutes. In a recent paper, investigators have shown up to 40 different fluorochromes can be used simultaneously to detect 40 different surface antigens [1, 2] from one sample. In conventional flow cytometry, cells pass through a very fine stream of liquid (generally PBS or equivalent). Depending on the experimental goal, cells are stained with antibodies that are conjugated directly with fluorochromes, like PE, have fluorescent proteins, like mCherry, or are stained by fluorescent dyes, like CFSE. Typically, cytometers have multiple lasers which are spatially separated. Fluorescent molecules can be excited by the lasers and emit fluorescence. Each fluorescent molecule has a very specific fluorochrome spectra which is unique to it.

Commercial flow cytometers collect fluorescence up to 800 nm (end of visible spectrum). A conventional system can have between 2 and 8 fluorescence light collection devices (detectors) dedicated to each laser. Different type of detectors can be used in flow cytometers. Detectors like Photo Multiplier Tubes (PMTs) have been a stable choice for many manufacturers over many years. These detectors are highly efficient at converting the light/incoming photons into photo electrons for detection. Later, acquisition software records the fluorescence in terms of number of photo electrons detected. New types of detectors, such as Avalanche Photo Diodes (APDs) and Silicon Photo Multipliers (SiPMs), are now gaining popularity among manufactures, as they are as good as PMTs, but more efficient in converting longer wavelength photons than PMTs [3, 4]. The photo electron generation process can be controlled by ‘PMT voltage’ for PMTs or by gain for APDs and SiPMs. Increasing PMT voltage or gain increases the detectors sensitivity but can also increase detector noise if increased too much, which negatively impacts data resolution [5].

In conventional systems, a small part of the fluorescence emission (generally part of the emission maxima) is chosen as the representative of the entire spectrum. A type of optical filter called band pass (BP) is used to choose which part of the emission need to be collected. These typically collect anything between 10 to 30 nm of the spectrum. For one fluorescence spectra, one detector acts as the main/primary detector and collects the light from the maximum emission region but other detectors (secondary), also collect part of the emission. The extent of the unwanted signal depends on the characteristics of the BP associated with the secondary detector and the shape/nature of the emission spectrum. We use a method called compensation to rectify the presence of the unwanted/extra signal in the secondary detectors. The details and rules of compensation have been extensively explained by others [6 – 10]. This unwanted signal is spillover. Spillover reduces the sensitivity, the lowest signal that can be detected over accumulation of signals coming from all other sources. In Flow Cytometers, spillover causes a non-linear spreading of properly compensated data (appears as straight line in log-log plot) [6]. Higher spillover causes higher spread in the secondary detector, which can mask dim signals. For immunophenotypic panels, spreading can decrease the detection sensitivity for low/dimly expressing antigens. The best way to avoid this problem is to find a combination of fluorochromes which cause no to low spread and assign antigens accordingly. For the purpose of designing a good panel, it is key to know how much spread can be expected in advance.

Useful tools/methods have been developed to aid investigators in this process. The most well-known and widely used is the Spillover Spreading Matrix (SSM), which provides a metric for signal variance introduced by spillover fluorescence for every fluorochrome-detector combination in a given panel [11]. The SSM can provide valuable insight into which combinations of fluorochromes are expected to work well together in a panel. Each value of the SSM is called Intrinsic Spillover Spread (ISS). However, care should be taken when using this method. We have found that ISS values are dependent on the detector sensitivity (detector voltage/gain). If users decrease the ‘PMT Voltage’ or gain of all the detectors, then ISS values drop for all detectors. This phenomenon can be used by the researcher to ‘reduce spread’ by lowering the voltage/gain (instead of redesigning the panel) while lower sensitivity can lead to reduced resolution between positive and negative populations. The effect of detector voltage/gain is intrinsically present in SSM and cannot be removed. It is also possible to use SSM to compare spread between machines, but the process is tedious (11). The output of the detectors is displayed using a scale, which can vary depending on the vendor and model. Aria Fusion from BD has 262,144 scale units (the scale is also known as dynamic range), CytoFLEX from Beckman Coulter has 16,777,215 and Quanteon from ACEA has 10^7.2^ scale units respectively. In this article we have introduced a new method to quantify spread, named Spread Quantification Index (SQI). SQI is independent of detector voltage/gain as well as independent of the dynamic range of the detector. It is easy to implement; researchers only need to carefully perform compensation by following all the rules to get a reproducible SQI value.

## Results

### Concept of SQI

The SQI value is defined as the difference in median fluorescent intensity (MFI) between the 99^th^ percentile and the 50^th^ percentile in detector B, of the positive population of fluorochrome X (Fig 1). Detector B is the recipient detector where spread from fluorochrome X is being registered. This value is a measure of total spread including spillover fluorescence in a secondary detector. Some examples of SQI can be found in Fig 2. The following abbreviations are used to describe the calculation procedure as explained in Fig 1.

**Figure 1:**
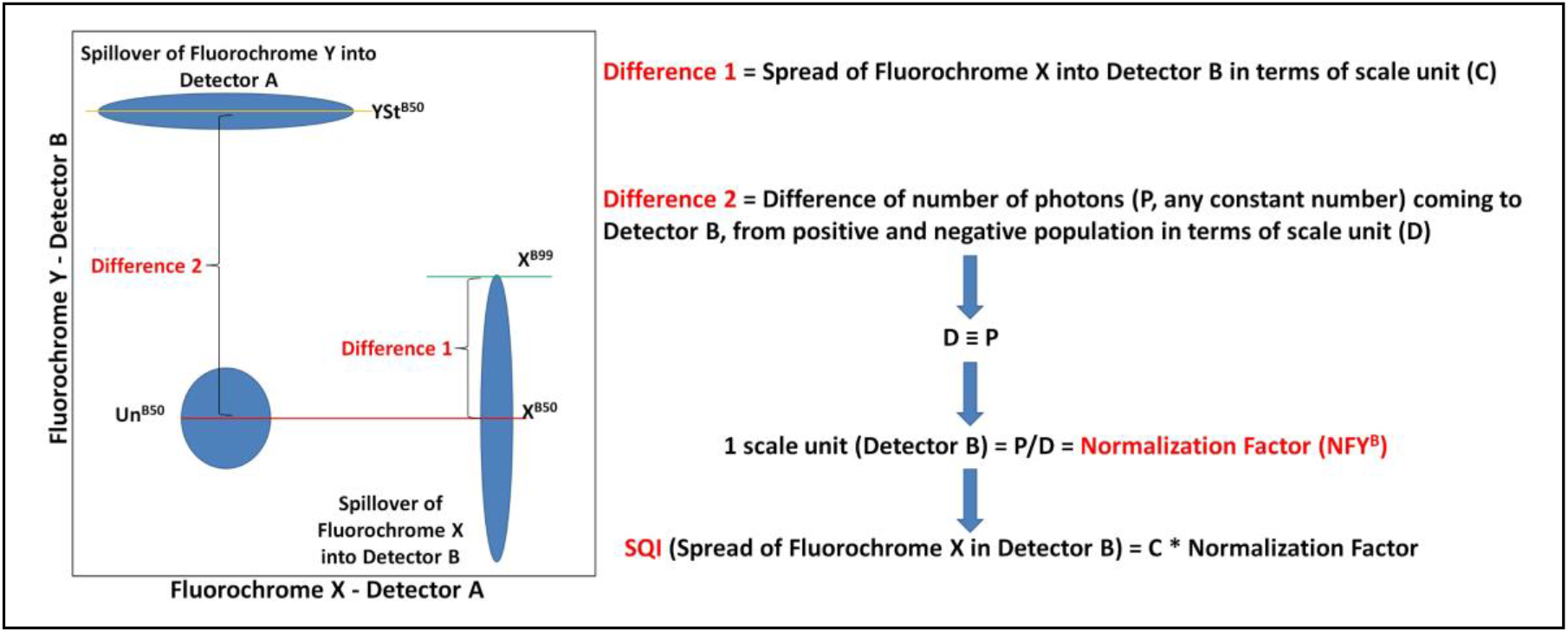
Concept of the SQI: The figure shows the concept of the SQI value. Spillover fluorescence from primary fluorochrome X results in spread in detector B, and vice versa. The spread is quantified as the difference of the 99th percentile and 50th percentile in dimension B. Subsequently, this value is multiplied by a normalization factor to make the SQI value independent of detector gain settings. The 99^th^ percentile is marked by a Green line.

**Figure 2:**
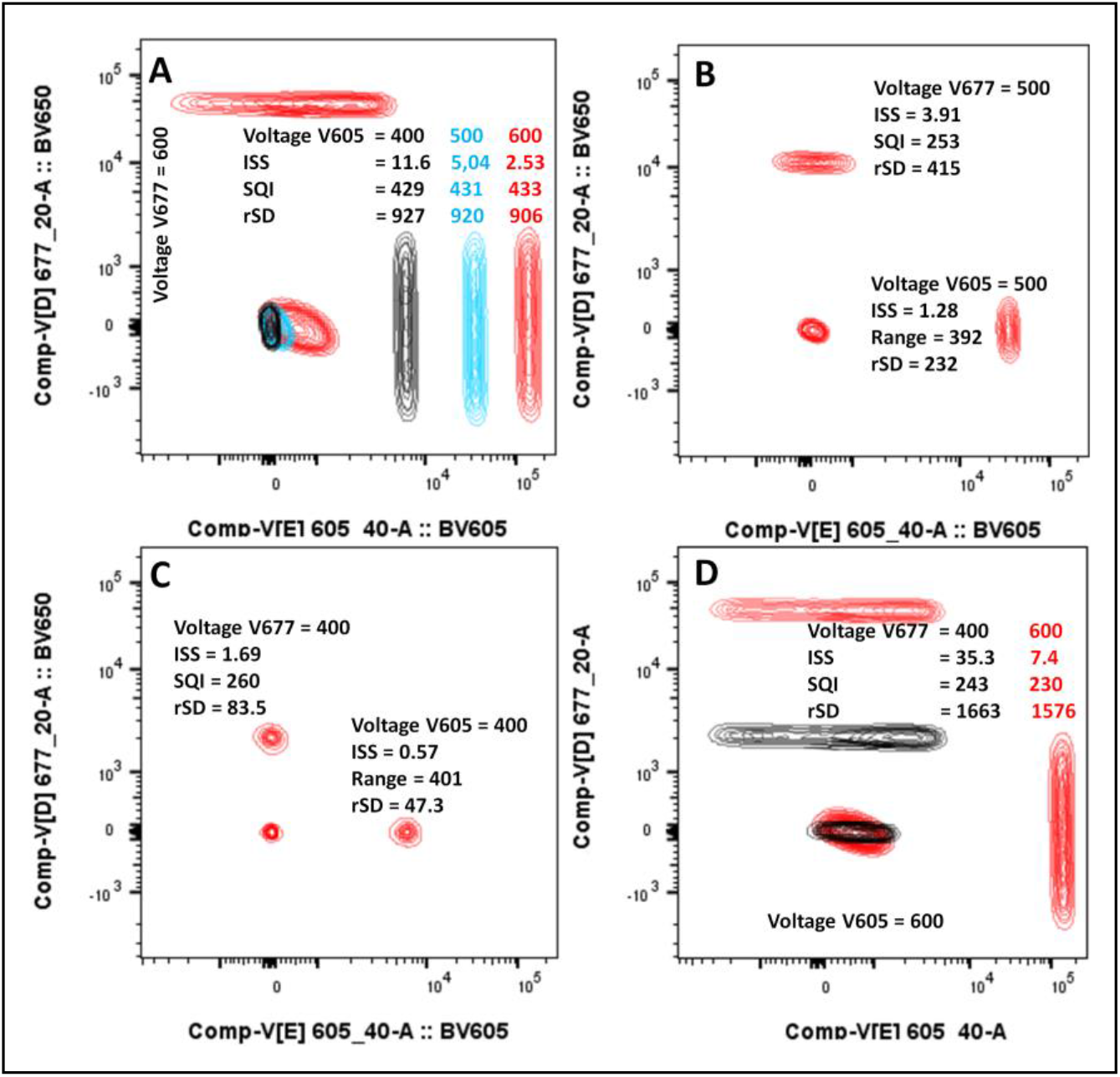
Effect of PMT voltage on ISS: In A, the primary detector (V605) voltage was changed from 600V to 400V in steps of 100V. This did not change the rSD and SQI values measuring the spread in the secondary detector (V677). However, the ISS values changed significantly. In A to C, voltages of both detectors were changed simultaneously to 600V, 500V or 400V respectively. The SQI values remained the same for both detectors in all cases, but the ISS values decreased considerably. In D only voltage of V677 was lowered which resulted in a huge increase of ISS but SQI remain same.

X^B50^ = 50^th^ percentile of compensated positive population of fluorochrome X in detector B

X^B99^ = 99^th^ percentile of compensated positive population of fluorochrome X in detector B

Un^B50^ = 50^th^ percentile of compensated unstained population of in detector B

YSt^B50^ = 50^th^ percentile of compensated stained population of fluorochrome Y in detector B

NFY^B^ = Normalization factor of fluorochrome Y in detector B (Normalization Factor, in Fig 1) (YSt^B50^ - Un^B50^) = The difference between the positive and negative populations (scale units) (Difference 2, D in Fig 1).

If P (any constant number, in this article we have used P = 9000) is the number of photons received by the detector B and (YSt^B50^ - Un^B50^) are the number of scale units of detector B that represent P. Then (YSt^B50^ - Un^B50^) ≡ P, but not equal.

1 scale unit in detector B = P/(YSt^B50^ - Un^B50^) = NFY^B^

If SQI value X^B^ = Spread of fluorochrome X in detector B,

Then SQI X^B^ = (X^B99^ – X^B50^) * NFY^B^

By applying the stated normalization factor, the SQI value is made independent from voltage/gain settings of the detectors, and dynamic range. The normalization factor is founded on two assumptions. First, the number of incoming photons remains the same when settings of the secondary detector are changed. This means that changing the detector voltage/gain will not change the excitation of the fluorochrome by the laser and the number of emitting photons will remain same when they pass through the same filters and mirrors. Second, differences in signal intensities are linearly proportional to changes in detector settings. This means that the conversion of photons into photo electrons is constant. In one gain setting, 1 photon converts into 3 photoelectron and 5 photons into 15 photoelectrons, while in another gain setting, if 1 photon converts to 5 photoelectrons, then 5 photons must convert into 25 photoelectrons.

The normalization factor allows for the SQI value to be determined without prior optimization of detector settings, provided that all obtained data is within the linear range and negative values are outside the detector noise. P is the difference in number of photons that are coming from positive and negative population in detector B. This number will stay practically constant when saturatedly stained samples will be maximally excited. The lasers of almost all of the presently available cytometers are designed to get maximal excitation. For this reason, we can get the same P values in all commercial cytometers. Maximal excitation makes P practically a constant. The constant nature of P in these terms allows for direct comparison of SQI values between instruments with different dynamic ranges. SQI showed a high degree of reproducibility in an average of three runs on the Fusion (Supplementary Figure S1). In the discussion, we elaborate on the conditions that must be met. Importantly, when multiple fluorochromes are associated with the same detector, the same amount of spread results in a higher SQI in secondary detectors when associated with dim fluorochromes (Supplementary Table S1).

### Classification of SQI values

Since there are no clear consensus or characteristics that divide spread in categories [11, 12], we created a different, semi-quantitative method to categorize SQI outcome as described below: Percentage of channels occupied by spread = [(YSt^B50^ - Un^B50^)/ total channel number] * 100 For example, 262,144 is the total number of scale units for BD machines (Fusion and Symphony). SQI between 0 to 120, 121 to 220, 221 to 300 and 300+ matches nicely with 0 to 0.5%, 0.5% to 1%, 1% to 1.5% and 1.5%+ of scale unit occupancy respectively. Comparing percent of scale units occupied with SQI will provide the user with a better understanding of spread between different types of machines (Table 1) and between multiple machines of the same type (Table 2). We have color coded SQI values in 4 categories (Table 1 and 2): 1 – 120 (Green); 121 – 220 (Yellow); 221 – 300 (Orange); and 300+ (Red). While Green and yellow both provide acceptable discriminative power in most cases, Green should be reserved for markers with low or unknown expression levels. Values with an Orange or Red SQI value should be used for markers on distinctly separated cell populations or for markers with exclusive expression on specific cell subsets. For example, if BV421 is combined with Alexa Fluor 647 (AF647), the SQI value is 20 (Green). Note that the SQI value is not zero due to measurement error of the autofluorescent signal and biological variance. In comparison, if PE-Cy5 is combined with AF647 the SQI value is 611 (Red). At these different levels of spread, 15% and 5% double positive events were identified respectively (Supplementary Fig S2). Double positive events are those which are outside the spreading error, received by the secondary/recipient detector (R670). The lower number of double positive events registered for the PE Cy5 compare to BV421 because of the need to account the higher spreading error coming from the PE Cy5.

**Table 1:** Comparison of spread in four different flow cytometers: Ten different single stained bead sets were run on four different instruments. SQI was calculated for every combination. For BV711 there was no data available for the Cytoflex S. The SQI values are categorized as follows: 1 – 120 (Green); 121 – 199 (Yellow); 200 – 299 (Orange); and 300+ (Red).

|  | Detector | BB700 | BV421 | BV605 | BV711 | PE | PE-CF594 | PE-Cy5 | PE-Cy7 | APC | APC-R700 |
| --- | --- | --- | --- | --- | --- | --- | --- | --- | --- | --- | --- |
| System | Fluorochrome | SQI | SQI | SQI | SQI | SQI | SQI | SQI | SQI | SQI | SQI |
| Symphony | BB700 |  | 45 | 68 | 563 | 23 | 18 | 38 | 56 | 523 | 509 |
| Fusion |  |  | 29 | 158 | 1819 | 8 | 17 | 43 | 74 | 329 | 471 |
| Cytoflex |  |  | 20 | 106 |  | 5 | 6 | 51 | 38 | 181 | 567 |
| Quanteon |  |  | 54 | 228 | 827 | 30 | 33 | 37 | 55 | 124 | 279 |
| Symphony | BV421 | 13 |  | 33 | 27 | 22 | 15 | 14 | 9 | 35 | 20 |
| Fusion |  | 13 |  | 44 | 26 | 7 | 10 | 10 | 11 | 23 | 17 |
| Cytoflex |  | 7 |  | 38 |  | 4 | 3 | 5 | 5 | 38 | 20 |
| Quanteon |  | 33 |  | 46 | 35 | 29 | 30 | 26 | 25 | 37 | 38 |
| Symphony | BV605 | 73 | 39 |  | 177 | 129 | 220 | 81 | 52 | 83 | 77 |
| Fusion |  | 110 | 92 |  | 220 | 61 | 248 | 87 | 61 | 89 | 107 |
| Cytoflex |  | 85 | 70 |  |  | 44 | 115 | 29 | 29 | 47 | 35 |
| Quanteon |  | 95 | 68 |  | 145 | 103 | 145 | 77 | 46 | 62 | 78 |
| Symphony | BV711 | 85 | 48 | 23 |  | 22 | 15 | 17 | 41 | 161 | 604 |
| Fusion |  | 75 | 85 | 32 |  | 7 | 9 | 15 | 62 | 92 | 476 |
| Cytoflex |  |  |  |  |  |  |  |  |  |  |  |
| Quanteon |  | 97 | 79 | 40 |  | 29 | 29 | 28 | 54 | 62 | 374 |
| Symphony | PE | 105 | 16 | 128 | 74 |  | 260 | 97 | 49 | 105 | 31 |
| Fusion |  | 125 | 17 | 334 | 101 |  | 197 | 96 | 55 | 109 | 39 |
| Cytoflex |  | 71 | 20 | 277 |  |  | 88 | 32 | 27 | 43 | 43 |
| Quanteon |  | 91 | 47 | 182 | 53 |  | 153 | 71 | 38 | 58 | 44 |
| Symphony | PE-CF594 | 215 | 18 | 184 | 150 | 156 |  | 178 | 103 | 198 | 51 |
| Fusion |  | 309 | 19 | 656 | 254 | 67 |  | 183 | 123 | 208 | 68 |
| Cytoflex |  | 201 | 19 | 480 |  | 42 |  | 49 | 66 | 64 | 103 |
| Quanteon |  | 202 | 47 | 334 | 93 | 124 |  | 134 | 73 | 94 | 70 |
| Symphony | PE-Cy5 | 490 | 22 | 28 | 305 | 61 | 41 |  | 192 | 855 | 319 |
| Fusion |  | 736 | 29 | 51 | 629 | 83 | 32 |  | 232 | 935 | 343 |
| Cytoflex |  | 649 | 21 | 39 |  | 34 | 13 |  | 156 | 816 | 453 |
| Quanteon |  | 435 | 49 | 51 | 370 | 127 | 47 |  | 141 | 326 | 201 |
| Symphony | PE-Cy7 | 31 | 16 | 21 | 29 | 43 | 26 | 18 |  | 36 | 41 |
| Fusion |  | 23 | 20 | 30 | 27 | 203 | 23 | 46 |  | 20 | 36 |
| Cytoflex |  | 22 | 20 | 23 |  | 306 | 7 | 131 |  | 41 | 25 |
| Quanteon |  | 36 | 47 | 39 | 46 | 74 | 37 | 30 |  | 38 | 47 |
| Symphony | APC | 105 | 18 | 20 | 120 | 22 | 18 | 128 | 52 |  | 280 |
| Fusion |  | 121 | 20 | 29 | 145 | 8 | 12 | 136 | 56 |  | 160 |
| Cytoflex |  | 111 | 22 | 24 |  | 5 | 10 | 145 | 32 |  | 264 |
| Quanteon |  | 121 | 48 | 39 | 127 | 32 | 33 | 129 | 49 |  | 165 |
| Symphony | APC-R700 | 51 | 16 | 20 | 117 | 22 | 14 | 35 | 85 | 366 |  |
| Fusion |  | 42 | 17 | 26 | 219 | 7 | 10 | 34 | 114 | 570 |  |
| Cytoflex |  | 105 | 21 | 22 |  | 5 | 2 | 83 | 60 | 179 |  |
| Quanteon |  | 59 | 48 | 36 | 173 | 30 | 29 | 40 | 88 | 267 |  |

**Table 2:** Comparison of all four Fusion setup: The SQI values for 10 fluorochromes run on three different Fusions are shown. For Fusion 2, “2_High” denotes higher laser power as for the other two instruments. The SQI values are categorized as follows: 1 – 120 (Green); 121 – 199 (Yellow); 200 – 299 (Orange); and 300+ (Red).

|  | Detector | B695 | V450 | V610 | V710 | Y582 | Y610 | Y670 | Y780 | R670 | R730 |
| --- | --- | --- | --- | --- | --- | --- | --- | --- | --- | --- | --- |
| Fusion | Fluorochrome | SQL | SQL | SQL | SQL | SQL | SQL | SQL | SQL | SQL | SQL |
| 1 | BB700 |  | 29 | 158 | 1819 | 8 | 17 | 43 | 74 | 329 | 471 |
| 2 |  |  | 28 | 105 | 1248 | 6 | 14 | 40 | 63 | 266 | 420 |
| 3 |  |  | 50 | 77 | 525 | 7 | 15 | 41 | 60 | 205 | 641 |
| 2_High |  |  | 29 | 107 | 1280 | 6 | 15 | 40 | 63 | 229 | 375 |
| 1 | BV421 | 13 |  | 44 | 26 | 7 | 10 | 10 | 11 | 23 | 17 |
| 2 |  | 16 |  | 42 | 23 | 6 | 8 | 11 | 13 | 20 | 17 |
| 3 |  | 14 |  | 35 | 20 | 6 | 9 | 10 | 10 | 15 | 13 |
| 2_High |  | 15 |  | 42 | 24 | 7 | 10 | 13 | 15 | 20 | 17 |
| 1 | BV605 | 110 | 92 |  | 220 | 61 | 248 | 87 | 61 | 89 | 107 |
| 2 |  | 110 | 77 |  | 177 | 40 | 194 | 86 | 57 | 79 | 102 |
| 3 |  | 101 | 43 |  | 175 | 42 | 165 | 89 | 54 | 79 | 89 |
| 2_High |  | 119 | 53 |  | 183 | 40 | 198 | 90 | 60 | 83 | 99 |
| 1 | BV711 | 75 | 85 | 32 |  | 7 | 9 | 15 | 62 | 92 | 476 |
| 2 |  | 84 | 91 | 27 |  | 6 | 8 | 16 | 55 | 91 | 586 |
| 3 |  | 84 | 55 | 24 |  | 6 | 8 | 16 | 53 | 63 | 709 |
| 2_High |  | 79 | 73 | 28 |  | 6 | 8 | 17 | 55 | 89 | 538 |
| 1 | PE | 125 | 17 | 334 | 101 |  | 197 | 96 | 55 | 109 | 39 |
| 2 |  | 121 | 15 | 400 | 81 |  | 163 | 99 | 55 | 98 | 46 |
| 3 |  | 130 | 15 | 148 | 86 |  | 172 | 100 | 50 | 92 | 31 |
| 2_High |  | 115 | 15 | 396 | 77 |  | 164 | 104 | 56 | 103 | 46 |
| 1 | PE-CF594 | 309 | 19 | 656 | 254 | 67 |  | 183 | 123 | 208 | 68 |
| 2 |  | 245 | 17 | 744 | 148 | 51 |  | 184 | 116 | 183 | 78 |
| 3 |  | 272 | 19 | 239 | 167 | 54 |  | 188 | 111 | 177 | 55 |
| 2_High |  | 241 | 17 | 747 | 141 | 53 |  | 193 | 118 | 191 | 77 |
| 1 | PE-Cy5 | 736 | 29 | 51 | 629 | 83 | 32 |  | 232 | 935 | 343 |
| 2 |  | 494 | 19 | 35 | 394 | 44 | 26 |  | 204 | 564 | 321 |
| 3 |  | 574 | 26 | 33 | 339 | 42 | 27 |  | 204 | 548 | 225 |
| 2_High |  | 499 | 19 | 35 | 418 | 34 | 26 |  | 212 | 611 | 308 |
| 1 | PE-Cy7 | 23 | 20 | 30 | 27 | 203 | 23 | 46 |  | 20 | 36 |
| 2 |  | 26 | 18 | 28 | 24 | 159 | 18 | 39 |  | 23 | 28 |
| 3 |  | 22 | 14 | 23 | 22 | 34 | 18 | 16 |  | 18 | 25 |
| 2_High |  | 21 | 17 | 28 | 25 | 57 | 18 | 20 |  | 23 | 25 |
| 1 | APC | 121 | 20 | 29 | 145 | 8 | 12 | 136 | 56 |  | 160 |
| 2 |  | 95 | 17 | 26 | 100 | 6 | 10 | 110 | 51 |  | 163 |
| 3 |  | 116 | 15 | 22 | 119 | 7 | 10 | 118 | 49 |  | 147 |
| 2_High |  | 110 | 18 | 27 | 116 | 6 | 11 | 113 | 51 |  | 139 |
| 1 | APC-R700 | 42 | 17 | 26 | 219 | 7 | 10 | 34 | 114 | 570 |  |
| 2 |  | 43 | 15 | 24 | 204 | 6 | 8 | 30 | 90 | 224 |  |
| 3 |  | 48 | 14 | 21 | 123 | 6 | 8 | 31 | 85 | 105 |  |
| 2_High |  | 42 | 15 | 25 | 212 | 6 | 8 | 31 | 90 | 104 |  |

### Intrinsic Spillover Spread values are dependent on detector voltage/gain settings

The ISS value of BV605 changed in the recipient detector when voltage of the primary detector changed, because the intensity difference (ΔF) between the positive and negative populations incorrectly describes a change in photonic input in the secondary detector (Fig 2A). Importantly, the variance as represented by the rSD and SQI value remained the same. Importantly, when voltages of both the detector were reduced, the rSD and ISS values were lower too (Fig 2A - C). Conversely, the SQI value was unchanged. In Fig 2D we only reduced the PMT voltage of the secondary detector which decreased the ΔF of the secondary detector resulting in increasing the related ISS.

### SQI values are independent of detector gain settings

Our main goal is to quantify spread independent of detector voltage/gain. The normalization factor was introduced for this reason. We devised this simple experiment to test the accuracy of the normalization factor. Single stained beads were measured with two different PMT voltage sets. The voltage was manipulated to position the positive population to 110,000 MFI or 50,000 MFI in all detectors. The SQI values were the same in both voltage sets, but the ISS values were quite different (Table 3). This observation is possible because SQI does not include any component from the primary detector. When the voltage of detector B was decreased, the spread from fluorochrome X in detector B was reduced, but the intensity of fluorochrome Y, which was used to calculate normalization factor, reduced proportionally, rendering SQI values independent of detector voltage. In contrast, when the same data were used to obtain the conventional Spillover Spread Matrices, the lower voltage set showed lower ISS values (Table 3). In another scenario, a series of voltages were applied in which the voltage of the secondary detector was kept constant, but the voltage of the primary detector was changed gradually (Supplementary Figure S3 A, B). Alternatively, the voltage of the primary detector was kept constant and the voltage of the secondary detector was changed (Supplementary Figure S3 C,D). In both cases the SQI values remained the same, however, the ISS values changed considerably.

**Table 3.** PMT voltage independent measure of spread: Two different sets of PMT voltages were applied to set the MFI of the positive population to 110,000 (110K) or 50,000 (50K). Six single stained bead samples were run using these two voltage sets, and the ISS and SQI values were calculated. The SQI values were similar at the different voltages, whereas the ISS values were voltage different.

|  | Detector | B695 |  | R730 |  | V710 |  | V610 |  | Y670 |  | Y610 |  |
| --- | --- | --- | --- | --- | --- | --- | --- | --- | --- | --- | --- | --- | --- |
| Fluorochrome | Intensity | 110K | 50K | 110K | 50K | 110K | 50K | 110K | 50K | 110K | 50K | 110K | 50K |
| BB700 | SQI |  |  | 577 | 596 | 632 | 632 | 77 | 78 | 39 | 40 | 15 | 17 |
|  | ISS |  |  | 7.24 | 5.27 | 9.27 | 6.59 | 1.08 | 0.78 | 0.53 | 0.37 | 0.2 | 0.15 |
| APC-R700 | SQI | 49 | 52 |  |  | 147 | 155 | 21 | 22 | 30 | 32 | 8 | 8 |
|  | ISS | 0.66 | 0.44 |  |  | 2.09 | 1.44 | 0 | 0 | 0.4 | 0.28 | 0 | 0.04 |
| BV711 | SQI | 89 | 96 | 786 | 788 |  |  | 24 | 25 | 15 | 16 | 8 | 8 |
|  | ISS | 1.22 | 0.84 | 8.98 | 6.23 |  |  | 0.14 | 0.11 | 0.16 | 0.1 | 0.03 | 0 |
| BV605 | SQI | 106 | 112 | 78 | 80 | 177 | 181 |  |  | 85 | 86 | 206 | 212 |
|  | ISS | 1.49 | 1.03 | 1.14 | 0.79 | 2.44 | 1.67 |  |  | 1.21 | 0.82 | 3.05 | 2.16 |
| PE-Cy5 | SQI | 721 | 785 | 309 | 298 | 412 | 411 | 38 | 39 |  |  | 31 | 29 |
|  | ISS | 11.03 | 7.71 | 4.31 | 3.02 | 5.8 | 4.14 | 0.44 | 0.33 |  |  | 0.37 | 0.26 |
| PE-CF594 | SQI | 299 | 322 | 63 | 62 | 198 | 195 | 296 | 301 | 179 | 184 |  |  |
|  | ISS | 4.27 | 2.88 | 0.83 | 0.57 | 2.67 | 1.82 | 4.16 | 2.87 | 2.42 | 1.73 |  |  |

### Application of SQI: Compare instruments with different dynamic range

To investigate which instrument is the best for running a given panel in terms of spread, we compared the SQI values between a Fusion, Symphony, Quanteon and a Cytoflex (Table 1). The Cytoflex showed the lowest number of SQI values above 120. The other instruments showed a higher number of values above 120. The Quanteon and Cytoflex had the lowest number of SQI values over 300. The Quanteon had the highest number of SQI values between 120 to 199.

### Application of SQI: Compare instrument performance

While all three Fusions had similar optical configuration and laser power, Fusion 3 performed best with only 6 instances of 300+ range. The performance of The Fusion 1 was considered worst with 12 occurrences of Range values over 300. The findings are summarized in Table 2. The results suggest that Fusion 3 is more suitable for the most sensitive experiments.

### Application of SQI: Investigate the effect of instrument characteristics on spread

SQI was used to investigate the effect of high and low laser powers in Fusion 2 which showed some interesting findings. At an output power of 140mW, the Red laser resulted in lower spillover spread from APC and APC-R700 compared to lower output power of 100mW (Table 2). In other cases, higher laser power showed minimal effect on spread. For instance, at 100mW, the 405nm laser showed virtually the same spread in all three violet detectors as compared to measurement at 85mW (Supplementary Table 2). Our data indicates that in our systems, higher laser powers other than for the red laser, doesn’t lower the data spread (Table 2).

### Validation assay

To test the usefulness of SQI, we ran two small panels separately. Antibodies in both panels have identical antibody clones and the exact same amount of antibody was used. In Panel 2, we replaced CD3-BV605, CD45-Alexa Fluor 488 with CD3-APC-R700 and CD45-BV650. We did this change to introduce more spread in the APC and PerCP-eFluor 710 detectors. We also choose BUV395, a dim fluorochrome with CD25 (low expression) to show clearly the adverse effect of lowering the gain. Both panels were run four times separately using four different gain settings (details are in material and methods). When analyzing for specific populations (Supplementary Figure S4), ISS and SQI calculated to show with change of gain, SQI remains the same but ISS drops (Supplementary Figure S5). Further, reducing gain resulted in the loss of data resolution of the specific populations, marked by the black arrows in panel 1, and spread-induced masking of the specific populations, marked by the red arrows in panel 2 (Supplementary Figure S5). These data, combined with the single color SQI with these same antibodies (Supplementary Table S4), demonstrate that BV650 and APC-R700 would produce significant spread in APC and PerCP-eFluor 710 detectors resulting in masking of dimly expressing cells.

## Discussion

To show how ISS values are dependent on detector voltage/gain, we explored two scenarios. First, the voltage of the primary detector was changed (Fig 2A). The spread remained the same as was evident by the unchanged rSD, but the ISS values changed because the value of √ΔF changed. More specifically, the lower voltage on the primary detector increased the ISS value, because the same Δσ_C_ [Equation 1 of reference 11] was divided by a smaller number (smaller ΔF). In the Supplementary Fig S3, we have shown SQI remains unchanged if we change primary or secondary voltages. However, the voltage changes do affect the ISS values considerably. In the second scenario (Fig 2A – C), voltages of the secondary detectors were also changed. In this example, lower voltages resulted in lower ISS values. Unfortunately, users have “optimized” the PMT voltages to get low ISS values. The motivation behind this is to reduce the spread so that they will be able to identify the double positive population better. This, however, may result in loss of resolution and obscuration of cells that have low expression of markers. Rather. users should adjust their panel to combinations that have low spread, increasing the chances of getting a good separation. Optimization of detector voltage/gain should be achieved by monitoring the separation of signal over the detector noise [5, 13]. SQI is not designed nor recommended for voltage/gain optimization. Because the SQI value is independent of detector voltage/gain, bias induced by voltage is prevented. The inherit relation of ISS and detector voltage/gain will always make the ISS values subjective. Even when users are not comparing different voltage sets, the influence of detector voltage/gain will always be there. SQI is, to the best of our knowledge, currently the only way to remove this subjectivity.

Furthermore, because variance and MFI are proportional to amplification settings in equal fashion, the MFI can be used to correct for the effect of changing secondary detector settings on spread. Additionally, the MFI of the unstained population (Un^B50^) is subtracted from the MFI of the Stained population (YSt^B50^) to normalize for differences in autofluorescence between channels. As shown in Table 3, lowering both voltages lowered the ISS values, with same samples As SQI is independent of voltage, the spread quantitation will remain the same. Ashhurst et al also showed that spread do not change with voltage alteration [10]. In Fig 2D we showed this by altering the PMT voltage of secondary detector which resulted in change of ISS but SQI remain same.

We presented a special scenario where the SQI value of one fluorochrome was calculated for two different fluorochromes, used separately in the secondary/recipient detector (Supplementary Table 1). For example, the SQI value of APC in detector B695 is different when different secondary fluorochromes (BB700 or PerCP-Cy^™^5.5) are considered. Because the SQI value is relative to the intensity of the secondary fluorochrome, the same spread from APC results in a relatively higher occupancy in channel number when combined with PerCP-Cy^™^5.5 as compared to BB700. It is helpful in the panel design phase to know that APC with PerCP-Cy^™^5.5 may be a more problematic combination for the identification of a double positive population. The calculation for ISS does not consider the MFI of the positive population from BB700 or PerCP-Cy^™^5.5. This makes the ISS values independent of the fluorochromes associated with the secondary detector. This will result in the spread in terms of ISS remains the same for both BB700 and PerCP-Cy^™^5.5. While this specific fluorochrome was not used in the validation panels, the concept can be seen in that panel. Specifically, when looking at CD3 vs TCRδγ, where the double positive population is much easier to identify with less spread in panel 1, when compared to panel 2. In addition, when gain is increased compared to the machine suggested values, there is a substantial increase in a single positive population that is not reported in the literature (Supplementary Figure S5). These interactions are also important to other fluorochromes within that staining panel, even with ideally designed panels. For example, even with panels designed to have little to no spread, gain values are still important, as decreasing the gain did not result in large changes in the above double positive population, but the data revealed a loss in a rare CD25+ CD127-population (Supplementary Figure S5), indicating that dim fluorochromes on antibodies for low expressing surface proteins are more sensitive to changes in detector sensitivity. The SQI and ISS values PE in BUV395 detector is not available for lower gain setups because the X^B99^ and X^B50^ values are outside the detector linear range which makes the calculation invalid. We deliberately introduced BV650 and APC-R700 in panel 2 to introduce spread in APC and PerCP-eFluor-710 detectors (Supplementary Table S4). In panel 1, CD3-CD4+ dim population lost the separation in lower gain value but because of the introduction of spread from BV650 and APC-R700 in APC channel becomes impossible to separate in panel 2. CD3+ TCRγδ ++ population is visible in both panels but due to the added spread CD3+ TCRγδ+ is not clearly visible in panel 2 (Red arrow) but in panel (Black arrow). We have found an interesting population in panel 1 (CD3-TCRγδ+) which gradually disappears with decrease in gain. To best of our knowledge, this is an unreported population. Because of spread introduced in panel 2, this population become unidentifiable. Importantly, these data (all SQI data for various machines in Tables 1-3 and Supplementary Table S4 and Figure S5) demonstrate that SQI is a platform independent method that will aid in better panel design.

It is not easy to compare ISS values between systems with different dynamic range, because instruments need to be calibrated first [11]. The ΔF accounts for the dependency of spread on the number of fluorescent photons, the ΔF needs to be numerically similar on both instruments to have a meaningful comparison of ISS values. This calibration is a tedious process. However, because the intensity values in the primary detector don’t influence the SQI values, this is not necessary in our method. Furthermore, due to the normalization factor, the scaling on every secondary detector is comparable. This makes the SQI value well suited to compare panel performance on different instruments.

As proof of concept, we compared the SQI values for a given set of single stains on different instruments with different dynamic range. Due to higher quantum efficiency for longer wavelength fluorescent light, we anticipated to find less spread in APD and SiPM based instruments in the red emission region as compared to PMT based instruments [3,4]. Indeed, both APD and SiPM detector-based instruments showed lower SQI values in detectors for APC and APC-R700 (Table 1). We matched the other instrument characteristics as best as possible, but these were not exactly the same. For example, laser output powers of the Quanteon and the Symphony were matched, but there was no option to alter the laser power for the Cytoflex. Furthermore, the filter configuration (characteristics and positioning) between the instruments was also not exactly the same (Supplementary Table S2). However, this example shows that it is possible to use SQI values for the purpose of comparing instrument performance with regards to spread for a given set of fluorochromes.

SQI values are empirical and very robust, because it’s not a variance-based calculation of spread. For a Gaussian distribution, the 99^th^ percentile can be calculated as 2.325 times the standard deviation (SD) of the mean. However, flow cytometric data follows the Poisson distribution and is known for positively skewed distributions [14]. For this reason, the factor of rSD required to calculate the 99^th^ percentile for each detector is highly variable. The median of 99^th^ percentile was obtained using FlowJo statistics tools. However, since the determination of the 99^th^ percentile is effectively based on just 1% of the data, it could potentially be distorted by outliers. To address this issue, careful singlet selection was performed, the median was used instead of the mean as a measure of central tendency, and a minimum of 35,000 single beads were recorded. This practice has been shown to provide highly robust results with less than 5% variation between tests (Supplementary Figure S1). While the SQI value is easy to use, there are some important things to keep in mind. First of all, the precision of the SQI value depends on correct ‘compensation’. For that reason, it is very important that all data is within the linear range of detection. Also, a thorough washing of the beads is an important criterion, as any non-specific binding of the antibody to the negative beads will change the photon difference (P) and introduce error in the normalization factor. Lastly, the SQI value is invalid if the correlation between variance and signal intensity is different at different amplification settings. Another important consideration to ensure reproducibility is the amount of bead bound antibody. This needs to be the same between repeats to ensure the difference in number of photons (P) between the positive and negative populations remain equal between runs. Using antibodies at saturating concentrations is therefore required. At the same time, this will also ensure maximum spread. On most of the commercially available machines, manufactures use a set of excitation optics to excite the fluorochromes maximally for practical use. The maximal excitation will ensure that almost the same number of photons will be emitted for both positive and negative populations. This is especially important with kits containing many different dyes, including tandem dyes. CD4 kits consist of many different tandem dyes. It has long been established that within different lots, the brightness and emission spectra of Tandem dyes differs [6,7]. To get reproducible data, it is required to use CD4 kits from same lot. This way, we can maintain the consistency of the emission spectra from the tandem dyes.

Nguyen *et al*. already proved that monitoring spread via ISS values is an excellent way to identify poor laser alignment, failing of the laser, other optical problems and to reveal detectors that may need optimization. As such, SQI can also be used to track systems over time. However, fluorochrome stability, of tandem dyes specifically, is of major concern, since tandem dye replacement or time dependent degradation will affect the outcome. A set of hard dyed plastic beads, which mimic single stains for all available channels would prove useful for long-term tracking.

## Conclusion

The main take home messages can be summarized as follows:

1. SQI values are unaffected by detector voltage/gain settings.
2. SQI values are independent of detector type or dynamic range. Therefore, SQI values can be directly compared between any instruments.
3. The SQI method is a useful tool to aid in panel design.

We strongly believe that this article helps novice users to understand data spread and how to deal with the effects better. Finally, we are certain that the voltage and dynamic range independent nature of SQI values will be instrumental in preventing manipulation of detector gain settings to minimize spread, rather than to use settings that provide the best separation and sensitivity.

## Materials and Methods

### Reagents

1. 5ml Falcon polystyrene round-bottom tubes (Corning Life Sciences, cat. no. 352054)
2. 5ml Falcon polystyrene round-bottom tube, with cell strainer snap cap (Corning Life Sciences, cat. no. 352235)
3. PBS as staining buffer
4. Human PBMC were isolated and stored as described in reference 15.

### Instruments

We have used 3 BD FACSAria™ Fusion cell sorters (Fusion) and a FACSSymphony A5 (Symphony) from BD Biosciences, a NovoCyte Quanteon from ACEA (Quanteon), and a CytoFLEX S (CytoFLEX) from Beckman Coulter Life Sciences. Fusion 1, 2 and 3 were used with identical optical configuration and laser output power unless stated otherwise. For one of the experiments, data was also generated at higher laser output powers on the Fusion 2 (Fusion 2_High). The output of all lasers of the Symphony and the Quanteon were set to 100mW. A detailed overview of the optical configurations (laser and filters) is given in the Supplementary Table S2. Description of Aurora can be found in reference [1].

### Antibody-fluorochrome conjugates

All experiments were done using human CD4 evaluation kit (BD Biosciences, cat no 566352). The following fluorochromes were used: BB700, PerCP-Cy™5.5, BV421, BV605, BV711, PE, PE-CF594, PE-Cy5, PE-Cy7, Alexa Fluor 647, APC, APC-R700 and Alexa Fluor 700. Clone matched (SK3) Alexa Fluor 647 conjugated CD4 antibody (cat no 344635), and anti-human Alexa Fluor 647 conjugated CD45RA antibody, clone HI100 (cat no 304153) were obtained from BioLegend. All reagents were stored as recommended by the manufacturer.

### Software

BD FACSDiva™ Software (version 8.0.1), NovoExpress, and CytExpert (version 2) were used only for data acquisition. Generation of compensation matrices and other analyses were done using FlowJo version 10.6.1.

### The Script

Compensated single stain fcs files are needed for the script to work. SQI values are computed using R (R 4.0.3). To do so, the population of stained beads, and unstained beads for fluorochrome X as well as the population of stained beads for fluorochrome Y need to be defined. This is done via statistical analysis using an analysis of mixture distribution. A first statistical model, using EM algorithm, is performed on the absolute value of the intensities measured by the primary detector corresponding to fluorochrome X. This analysis, performed with the R function normalmixEM from R package mixtools, decomposes the intensities measured on the primary detector as two distinct normal distributions, i.e., the unstained control beads and the stained beads populations. By taking the first and the third quartile of these distributions, low and high boundaries of both unstained and stained populations can be derived, which allows for gating of the data. The same approach is performed to gate the population of the fluorochrome Y stained beads. Once the three different population of beads are defined, the relevant parameters for SQI value computations are obtained. The script can be found in the webpage: github.com/RenTissier/RangeComputation.

### Data collection

At least 35,000 single beads or 100,000 single cells were recorded. We made sure that the population intensities stay within the linear dynamic range of the detector. Manufacturer recommended quality controls were performed before every experiment.

### Sample preparation

#### Staining of antibody capture beads

1. Antibody capture beads (both blank and positive) were vortex for ∼ 3 seconds.
2. One drop of each capture bead was mixed in a tube.
3. The saturating antibody concentration was determined in titration series for every antibody-fluorochrome conjugate used.
4. 0.25µg of antibody was added to all samples (at saturation). 0.25µg is well above the saturation amount for any antibody we have used.
5. The samples were stained at 4°C for 30 minutes in the dark.
6. Beads were washed twice in 3 ml of staining buffer, 5 minute centrifugation at 500 g (acceleration 9, deceleration 5).
7. Re-suspended in 200 µl of PBS.

#### Staining of PBMC

1. The cells were thawed in a 37°C water bath.
2. Filtered gently.
3. Centrifuged at 500 g for 10 minutes.
4. The supernatant was discarded.
5. 1 × 10^6^ cells were re-suspended in 100µl of staining buffer.
6. The saturating antibody concentration was determined in titration series for every antibody-fluorochrome conjugate used.
7. 0.06 µg of CD4 antibody was added (at saturation).
8. The samples were stained at 4°C for 30 minutes in the dark.
9. Beads were washed twice in 3 ml of staining buffer, 5 minute centrifugation at 500 g (acceleration 9, deceleration 5).
10. Re-suspended in 200 µl of staining buffer.

### Intrinsic Spillover Spread values are dependent on detector gain settings

Antibody capture beads were stained as mentioned above. Single stained beads were run only changing the primary PMT voltage from 600V to 400V in steps of 100V. A mixture of these single stains were run using different PMT voltages. ISS values were calculated and compared to the Robust Standard Deviation (rSD).

### SQI values are independent of detector gain settings

1. Saturated single stains of six different fluorochromes (BB700, BV605, BV711, PE-CF594, PE-Cy5 and APC-R700) were prepared as described above.
2. Two sets of PMT voltages were used.
3. Voltages in group one was set to place the positive population at ∼ 110,000 MFI.
4. Voltages in the second group were set to place the positive population at ∼ 50,000 MFI.
5. Range was calculated for all spillover signals.
6. In a separate experiment single stained beads were run for a series of PMT voltages. In one set only the voltage for the primary detector was changed, and in the second set only the voltage of secondary detector was changed. The ISS and SQI values were calculated for both sets.

### Application of SQI to compare instrument performance

We have three Fusions. The comparison was done using same laser power for all. Only in case of the Fusion 2 another higher laser power setup was used to test the effect of laser power on spread.

1. Single stains were run in all three Fusions.
2. Each experiment in each machine was performed in triplicates.
3. SQI value from all the repeat experiments were calculated and compared to check intra assay variation.

### SQI, as dynamic range independent measure of spread

1. Fusion, Symphony, Quanteon and Cytoflex were used for this comparison. Fusion and Symphony has a 5 decade scale. Cytoflex and Quanteon has 7 and 7.2 decade scale respectively.
2. For Quanteon and Cytoflex, positive signals were set to approximately 250,000 and 1,000,000 MFI. For the Symphony and Fusion these were set to 110,000 MFI. Spillover from BV711 was not analyzed in the Cytoflex, because it lacked the appropriate detector.
3. Saturated single stain antibody capture beads were run in each system in triplicates.
4. Range was calculated for all spillover signals.

### Application of SQI: Investigate the effect of instrument characteristics on spread

Single stained beads were run at different laser output powers. The SQI values were compared (same Fusion) to investigate the effect of excitation intensity on spread.

### Usefulness of SQI

1. Human PBMCs were used as samples
2. 7 × 10^6^ Cells were thawed in a 37°C water bath.
3. The cells were then filtered gently and centrifuged at 500 g for 10 minutes.
4. The supernatant was discarded, and the cells were re-suspended in 700 µl of staining buffer.
5. 100 µl of the cell suspension was kept aside as complete unstained.
6. 600 µl cells were stained 1^st^ with BV421 or PE-Cy5 or BB700 conjugated CD4 antibody (0.36 µg) for 15 minutes in dark at 4°C.
7. This was divided in 6 equal portions.
8. Five different amounts (0.01, 0.025, 0.05, 0.5 and 1.25 µg) of CD45RA antibody were added.
9. The samples were stained for another 30 minutes in dark at 4°C,
10. Cells were washed twice in 3 ml of staining buffer, 5 minute centrifugation at 500 g (acceleration 9, deceleration 5).
11. Re-suspended in 200 µl of staining buffer.
12. Single stains of BV421, PE-Cy5, BB700 and Alexa Fluor 647 were prepared using BD compensation beads as described previously for automated compensation.

### Staining for the validation assay

Cells were stained with different antibody as described above. Before adding the antibodies 5 µl of Brilliant stain buffer (BD, Catalog No 563794) and 5 µl True Stain Monocyte blocker (Biolegend, Catalog no 426101) was added in every tube. As recommended in reference [1], cells were 1^st^ stained with TCRγδ antibody for 15 minutes then stained with the master mix of the rest of the antibodies for another 15 minutes. Cells were then washed twice with 3 ml of staining buffer and finally resuspended in 250 µl of PBS, kept in ice until analyzed using a Cytek Aurora. Final staining volume was 200 µl. We have used two panels, 1 and 2. All the details are in Supplementary Table S3. One drop of compensation beads (UltraComp eBeads™ Compensation Beads: Thermo Fisher, catalog 01-2222-41) was stained with half of the antibody used to stain cell for 20 minutes and washed twice with PBS.

### Flow Cytometer setup for the validation assay

1. Both panels were run using exactly same gain settings.
2. Four sets of gain were used.
3. Cytek Assay Settings or CAS was used as standard gain setup.
4. One setup was created by increasing the gain of all fluorescence channel by 80%. Other two setups were created by decreasing the gain of all fluorescence channel by 50 and 70% respectively.
5. For every gain set antibody capture beads were run to perform the unmixing, followed by the fully stained cells.

## Data availability

All data are available at http://flowrepository.org, under the Repository ID of FR-FCM-Z336 to Z339 and FR-FCM-Z33A to Z33H

## Acknowledgement

The authors thank Dr. Heinz Jacobs for his critical comments. The authors also thank Dr. David Parks for his valuable remarks, comments and suggestions, especially with regards to the method of normalization. We like to thank Dr. Morten Nørgaard Andersen for his crucial critical comments. We thank Dr. Alexandra Terry for providing the cells. The authors thank Enver Delic to provide access to the CytoFLEX and BIOKE for access to the Quanteon. Martijn van Baalen helped DB tremendously to write the manuscript, gave critical inputs and generated/analyzed data for Supplementary Figure 3. DB like to thank Lyndsay Richard for all her help. This work is supported by institutional funding from the Dutch Cancer Society and grant from the National Institute on Aging, R00 AG055717 (MR).

## Author Contributions Statement

DB conceived, performed and analyzed experiments. DB and MR wrote the manuscript. RT developed the script. FvD actively helped in data collection and played a crucial role in instrument maintenance, which kept the flow cytometers in the best condition. AP helped in sample preparation.

## Supplementary Information

**Supplementary Table S1:** Effect of multiple fluorochrome in one detector on SQI: In this table we have presented a special scenario, where B695, R670 and R730 are associated with two fluorochromes. Data indicates same amount of spread (as channel number) translates in bigger SQI when a dim fluorochrome is associated with the secondary detector.

| Detector | B695 |  | V450 | V610 | V710 | Y582 | Y610 | Y670 | Y780 | R670 |  | R730 |  |
| --- | --- | --- | --- | --- | --- | --- | --- | --- | --- | --- | --- | --- | --- |
| Fluorochrome | BB700 | PerCP-Cy<sup>TM</sup>5.5 |  |  |  |  |  |  |  | Alexa 647 | APC | APC R700 | Alexa 700 |
| **BB700** |  |  | 29 | 161 | 1864 | 8 | 17 | 41 | 72 | 232 | 341 | 483 | 1695 |
| **PerCP-Cy<sup>TM</sup>5.5** |  |  | 20 | 26 | 258 | 8 | 9 | 56 | 63 | 53 | 78 | 117 | 410 |
| **Alexa Fluor 647** | 69 | 555 | 21 | 25 | 48 | 8 | 10 | 80 | 51 |  |  | 223 | 782 |
| **APC** | 119 | 954 | 20 | 30 | 147 | 8 | 12 | 134 | 55 |  |  | 162 | 568 |
| **APC-R700** | 42 | 335 | 17 | 26 | 223 | 7 | 9 | 35 | 116 | 386 | 569 |  |  |
| **Alexa Fluor 700** | 21 | 171 | 20 | 25 | 96 | 7 | 9 | 15 | 45 | 23 | 34 |  |  |

**Supplementary Table S2:** Configuration detail: This table lists the optical configurations of the instruments that were used in this study.

| Laser | BD FACSAria™ Fusion 1 and 3 |  |  | BD FACSAria™ Fusion 2 |  |  | BD FACSSymphony A5 |  |  | ACEA NovoCyte Quanteon |  |  | BC CytoFLEX S |  |
| --- | --- | --- | --- | --- | --- | --- | --- | --- | --- | --- | --- | --- | --- | --- |
|  | Long Pass | Band Pass | Laser Power (mW) | Long Pass | Band Pass | Laser Power (mW) | Long Pass | Band Pass | Laser Power (mW) | Mirror | Band Pass | Laser Power (mW) | Band Pass | Laser Power (mW) |
| 355 |  |  | NA | 770 | 820/60 | 50 | 770 | 819/44 | 100 |  |  | NA |  | NA |
|  |  |  |  | - | 379/28 |  | 690 | 735/50 |  |  |  |  |  |  |
|  |  |  |  |  |  |  | 630 | 670/25 |  |  |  |  |  |  |
|  |  |  |  |  |  |  | 600 | 615/20 |  |  |  |  |  |  |
|  |  |  |  |  |  |  | 550 | 580/20 |  |  |  |  |  |  |
|  |  |  |  |  |  |  | 450 | 515/30 |  |  |  |  |  |  |
|  |  |  |  |  |  |  | - | 379/28 |  |  |  |  |  |  |
| 405 | 750 | 780/60 | 85 | 750 | 780/60 | 100 | 750 | 780/60 | 100 | 757LP | 780/60 | 100 | 450/45 | 80 |
|  | 690 | 710/50 |  | 690 | 710/50 |  | 735 | 750/30 |  | - | 725/40 |  | 525/40 |  |
|  | 630 | 660/20 |  | 630 | 660/20 |  | 685 | 711/25 |  | 705LP | 695/40 |  | 610/20 |  |
|  | 595 | 610/20 |  | 595 | 610/20 |  | 635 | 661/11 |  | 685LP | 660/20 |  | 660/20 |  |
|  | 505 | 525/50 |  | 505 | 525/50 |  | 595 | 605/40 |  | 598SP | 615/20 |  |  |  |
|  | - | 450/50 |  | - | 450/50 |  | 550 | 586/15 |  | 570SP | 586/20 |  |  |  |
|  |  |  |  |  |  |  | 455 | 470/20 |  | 552LP | 530/30 |  |  |  |
|  |  |  |  |  |  |  | - | 431/28 |  | 495LP | 445/45 |  |  |  |
| 488 | 655 | 695/40 | 50 | 655 | 695/40 | 100 | 750 | 780/60 | 100 | 757LP | 780/60 | 100 | 690/50 | 50 |
|  | 502 | 530/30 |  | 502 | 530/30 |  | 735 | 750/30 |  | - | 725/40 |  | 525/40 |  |
|  | - | 488/10 |  | - | 488/10 |  | 685 | 710/50 |  | 705LP | 695/40 |  | 488/10 |  |
|  |  |  |  |  |  |  | 635 | 661/11 |  | 685LP | 660/20 |  |  |  |
|  |  |  |  |  |  |  | 600 | 610/20 |  | 598SP | 615/20 |  |  |  |
|  |  |  |  |  |  |  | 570 | 586/15 |  | 570SP | 586/20 |  |  |  |
|  |  |  |  |  |  |  | 505 | 530/30 |  | 552LP | 530/30 |  |  |  |
|  |  |  |  |  |  |  | - | 488/10 |  |  |  |  |  |  |
| 640 | 755 | 780/60 | 100 | 755 | 780/60 | 140 | 750 | 780/60 | 100 | 757LP | 780/60 | 100 | 660/20 | 50 |
|  | 690 | 730/45 |  | 690 | 730/45 |  | 690 | 730/45 |  | - | 725/40 |  | 712/25 |  |
|  | - | 670/30 |  | - | 670/30 |  | - | 670/30 |  | 705LP | 695/40 |  | 780/60 |  |
|  |  |  |  |  |  |  |  |  |  | 685LP | 660/20 |  | - |  |
| 561 | 735 | 780/60 | 50 | 735 | 780/60 | 50 | 750 | 780/60 | 100 | 757LP | 780/60 | 100 | 585/42 | 30 |
|  | 685 | 710/50 |  | 685 | 710/50 |  | 685 | 710/50 |  | - | 725/40 |  | 610/20 |  |
|  | 630 | 670/14 |  | 630 | 670/14 |  | 635 | 670/30 |  | 705LP | 695/40 |  | 690/50 |  |
|  | 600 | 610/20 |  | 600 | 610/20 |  | 600 | 610/20 |  | 685LP | 660/20 |  | 780/60 |  |
|  | - | 582/15 |  | - | 586/15 |  | - | 586/15 |  | 598SP | 615/20 |  |  |  |
|  |  |  |  |  |  |  |  |  |  | 570SP | 586/20 |  |  |  |
|  |  |  |  |  |  |  |  |  |  | 552LP | 561/14 |  |  |  |

**Supplementary Table S3:** Antibody details of the validation assay.

|  | Antibody | Fluorochrome | Clone | Catalog | Vendor | (ng of antibody/test) |  |
| --- | --- | --- | --- | --- | --- | --- | --- |
| Panel 1 | CD45 | Alx488 | HI30 | 304019 | Biolegend | 250 |  |
|  | CD3 | BV605 | UCHT1 | 300459 | Biolegend | 400 |  |
|  | CD123 | BV711 | 6H6 | 306030 | Biolegend | 125 | Panel 2 |
| | TCR$\gamma\delta$ | PerCP-eFluor™<br>710 | B1.1 | 46-9959-42 | Thermo | 500 | |
|  | CD4 | APC | SK3 | 566915 | BD | 100 |  |
|  | CD8 | BUV805 | SK1 | 612889 | BD | 125 |  |
|  | CD25 | BUV395 | 2A3 | 564034 | BD | 500 |  |
|  | CD127 | PE | HIL-7R-M21 | 557938 | BD | 500 |  |
|  | CD3 | APC-R700 | UCHT1 | 565120 | BD | 400 |  |
|  | CD45 | BV650 | HI30 | 304043 | Biolegend | 250 |  |

**Supplementary Table S4:** Use of SQI in panel design: Antibodies from Supplementary Table S3 used to saturatedly stain antibody capture beads to calculate SQI.

| SQI |  | Detector |  |  |  |  |  |
| --- | --- | --- | --- | --- | --- | --- | --- |
|  |  | APC | APC-R700 | Alexa Fluor 488 | BV605 | BV650 | PerCP-eFluor 710 |
| Fluorochrome | APC |  | 16 | 11 | 36 | 58 | 26 |
|  | APC-R700 | 104 |  | 16 | 24 | 45 | 523 |
|  | Alexa Fluor 488 | 23 | 10 |  | 6 | 18 | 4 |
|  | BV605 | 66 | 30 | 17 |  | 46 | 27 |
|  | BV650 | 231 | 42 | 21 | 141 |  | 51 |
|  | PerCP-eFluor 710 | 113 | 176 | 14 | 36 | 42 |  |

**Supplementary Figure S1:**
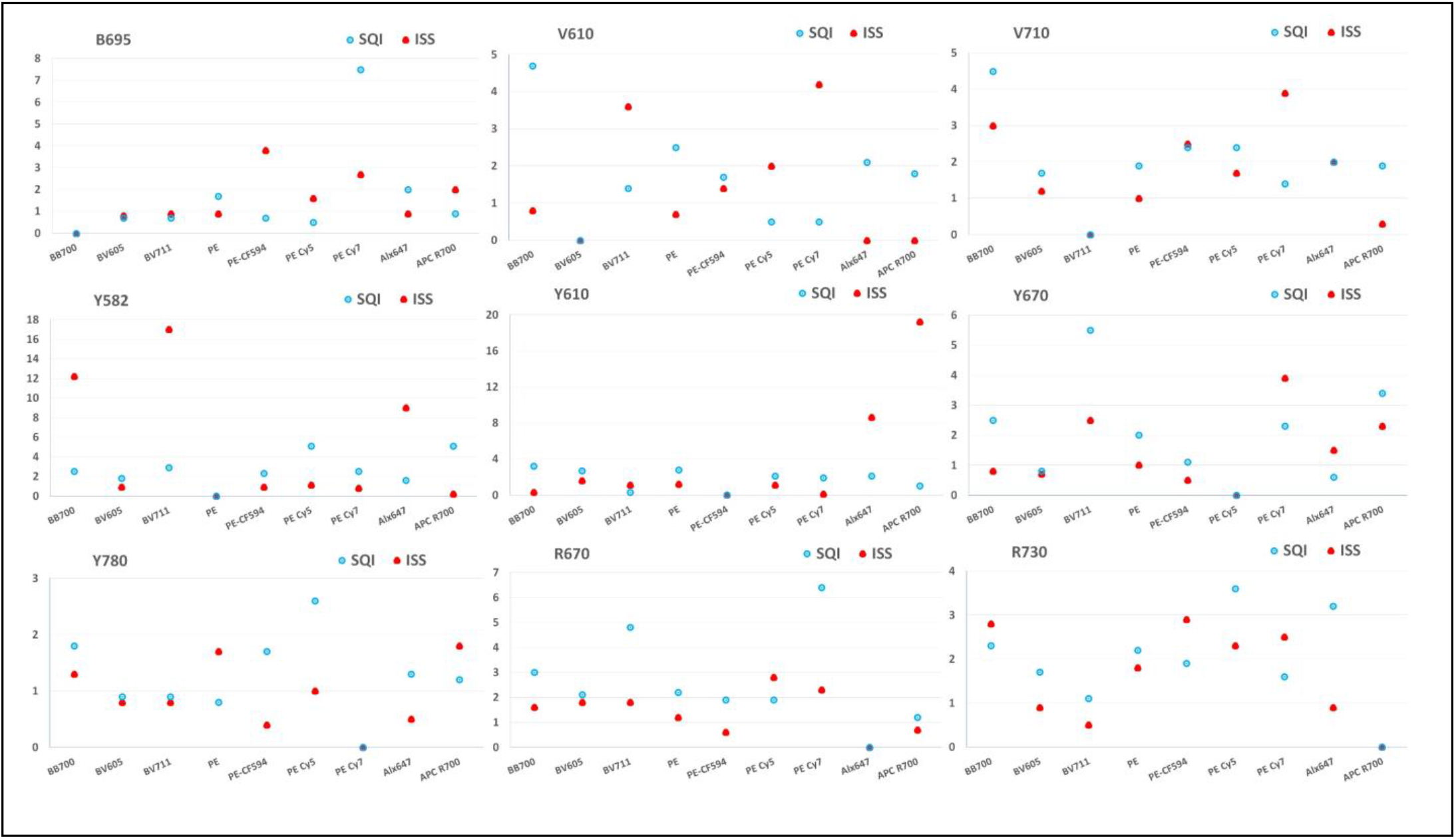
Reproducibility of SQI: Three replicative data sets from Fusion 1 were used to calculate SQI and ISS values. Each dot represents the relative standard deviation (%) of three runs. Blue circle and Red triangle represent SQI and ISS respectively.

**Supplementary Figure S2:**
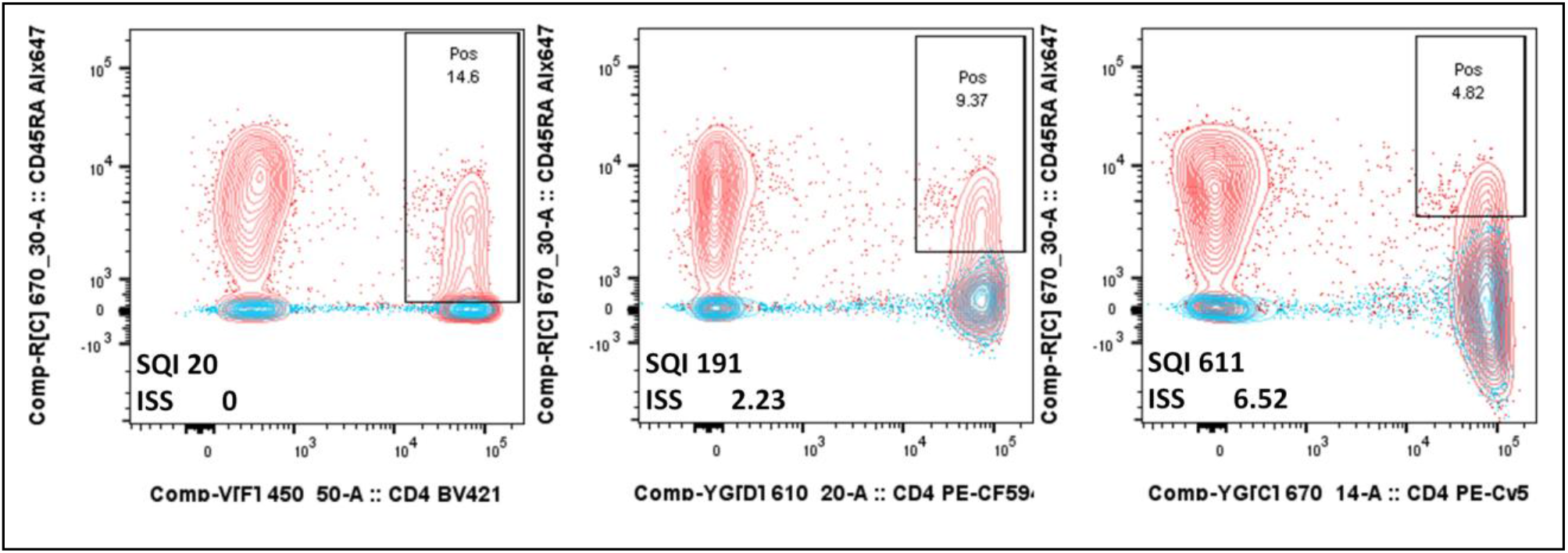
Usefulness of range: Alexa Fluor 647 conjugated CD45RA was combined with CD4 BV421, PE-CF594 or PE-Cy5 (clone matched), separately. The concentration of the CD45RA antibody was equal in all mixes. The blue contour indicates the control sample, which was stained with a CD4 antibody only. The red contour shows the combined staining. SQI and ISS values are displayed. The double positive population is better separated from the single positive population at lower SQI values as is exhibited by a higher percentage.

**Supplementary Figure S3:**
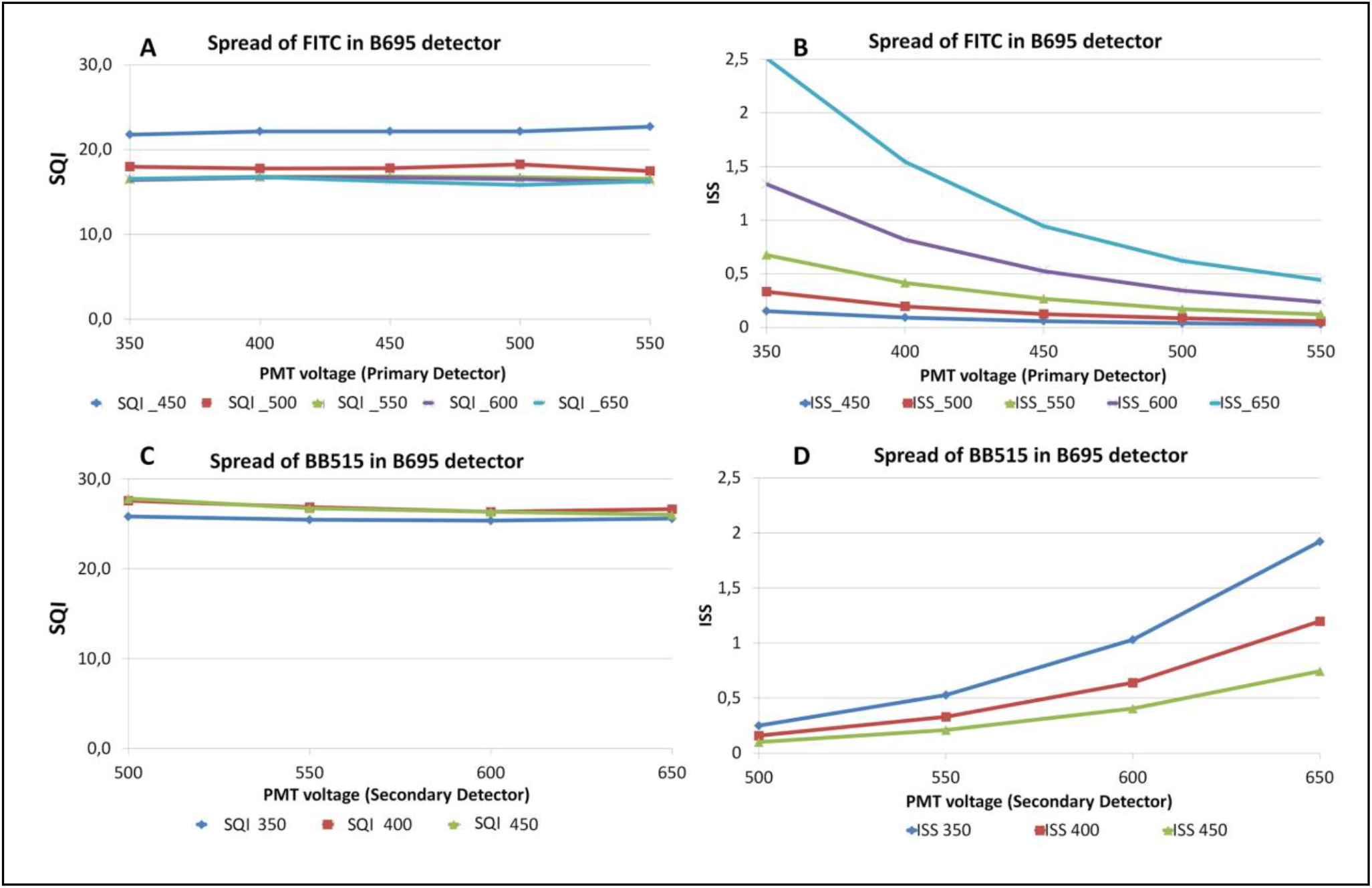
Effect of detector settings on SQI and ISS: In A and B we have only changed the PMT voltage for the primary detector. SQI remains same but ISS dropped with increase of voltage. In C and D, we only changed PMT voltage for secondary detector. SQI again remain same but ISS increased with voltage.

**Supplementary Figure S4:**
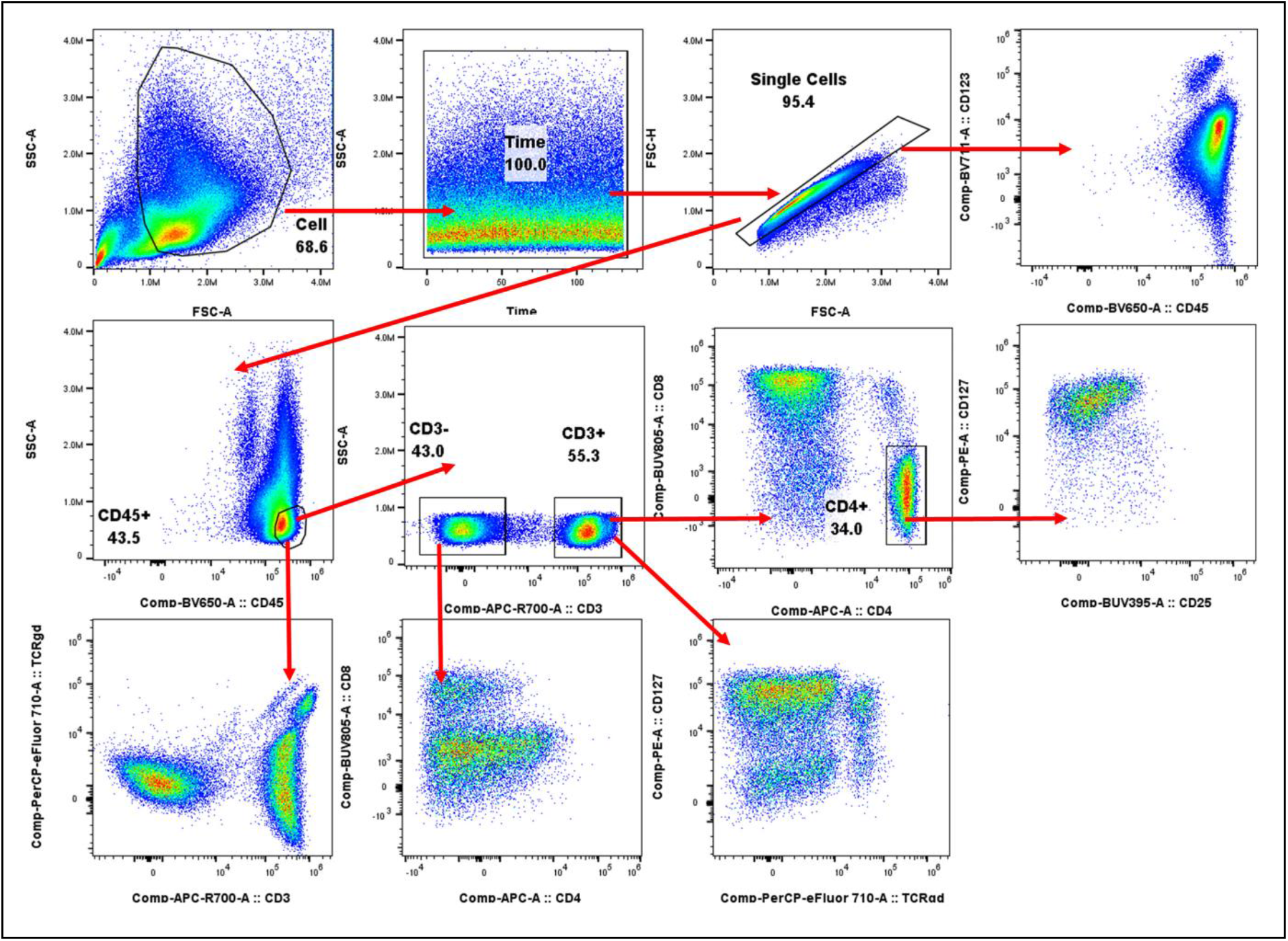
Analysis hierarchy of the validation assay. PBMCs were stained for flow cytometric analyses. Data were analyzed for the populations indicated.

**Supplementary Figure S5:**
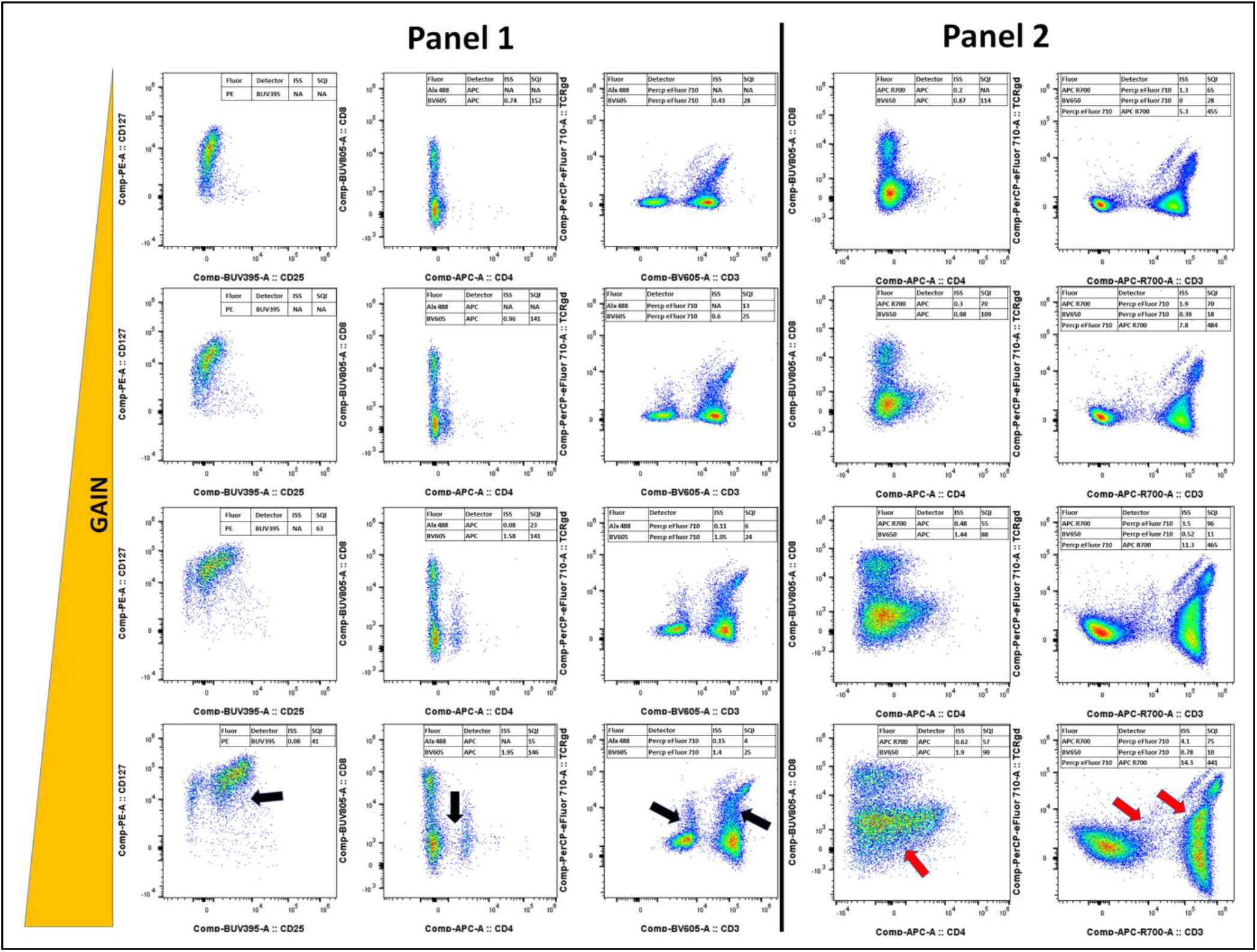
Changes to detector sensitivity artificially alters data presentation. Specific populations of PBMCs stained as in S4 are shown. Data were acquired on a Cytek Aurora, with standard gain values used (Cytek Assay Settings, CAS; 3^rd^ row across), with additional acquisition runs with gain in all channels increased by 80% (bottom row across) and gain in all channels decreased by 50% (2^nd^ row across) and 70% (Top row across). Panel 1 and Panel 2 differ in 2 fluorochrome labels, CD3-BV605, CD45-Alexa Fluor 488 and CD3-APC-R700, CD45-BV650, respectively. Black arrows indicate populations that lose resolution as the gain is decreased, and Red arrows indicate populations masked by spread and decreased gain. Numbers in each plot indicate the corresponding ISS and SQI values, where N/A is used when the MFI of the populations are out of the linear range of the detector.

